# IAP antagonists potentiate TNFα-triggered apoptosis but selectively eliminate senescent tumor cells independently of TNFα

**DOI:** 10.1101/2025.02.03.636172

**Authors:** Hiroaki Ochiiwa, Takeshi Wakasa, Yuki Kataoka, Koji Ando, Eiji Oki, Yoshihiko Maehara, Makoto Iimori, Hiroyuki Kitao

**Author notes:** **Corresponding authors:** Makoto Iimori, 2-15-1 Tamura, Sawara-ku, Fukuoka 814-0193, Japan., Hiroyuki Kitao, 2-15-1 Tamura, Sawara-ku, Fukuoka 814-0193, Japan.

## Abstract

Therapy-induced senescence (TIS) is a state of cell division arrest induced by chemotherapy that blocks tumor growth. TIS tumor cells affect the tumor microenvironment through their senescence-associated secretory phenotype and independently acquire stemness, which makes them more aggressive and causes relapse once they regrow. To eradicate tumors by chemotherapy, long-lived TIS tumor cells must be efficiently eliminated. Here, we show that AZD5582 and AT406, which are potent antagonists of inhibitor of apoptosis proteins (IAP antagonists) that suppress the activities of cellular inhibitor of apoptosis protein 1 (cIAP1), cellular inhibitor of apoptosis protein 2 (cIAP2), and X-linked inhibitor of apoptosis protein (XIAP), selectively induced apoptosis mediated by caspase 8 and effector caspases in TIS tumor cells, which produced and secreted tumor necrosis factor α (TNFα). However, these IAP antagonists were still selectively cytotoxic to TIS tumor cells even when TNFα was absent (TNFα-knockout cells) or neutralized (by a neutralizing antibody), indicating they have TNFα-independent senolytic activity. Consistently, these IAP antagonists also sensitized tumor cells that had been induced to become senescent by nutlin-3a, which activates p53 but does not trigger TNFα production. Furthermore, TNFα sensitized tumor cells treated with these IAP antagonists irrespective of their senescence status. Collectively, these data indicate that IAP antagonists that inhibit cIAPs and XIAP not only potentiate TNFα-triggered apoptosis but also have TNFα-independent senolytic activity. We propose that IAP antagonists are good concomitant drugs of chemotherapy that induces TIS, not only as senolytic drugs but also as sensitizers of adjacent non-senescent tumor cells mediated by paracrine TNFα.

## Introduction

Chemotherapy is one of the most important options for cancer treatment. However, it is difficult to eradicate all tumor cells *in vivo* because of intratumor heterogeneity, the unequal distribution of drugs, or different cell fate decisions according to the status of each tumor cell. To completely cure cancer, surviving tumor cells after chemotherapy must be eliminated. Tumor cells often enter a senescent state called therapy-induced senescence (TIS) when they stop growing due to chemotherapy^1^. TIS tumor cells tend to be resistant to chemotherapeutic drugs, which makes it difficult to completely eliminate tumor cells by continuous treatment with conventional therapy. The senescence-associated secretory phenotype (SASP) of TIS tumor cells also enforces the tumor microenvironment under chronic inflammation and mediates their patho-physiological effects^2^. Furthermore, cell-cycle re-entry often occurs under certain circumstances in TIS tumor cells, which acquire cancer stemness and tumor aggressiveness upon senescence^3,4^. Thus, although senescence is a potent tumor suppressive mechanism, the prolonged presence of TIS tumor cells poses significant clinical risks.

Senescent cells accumulate in various tissues and organs with aging and are thought to disrupt tissue structure and function mainly due to their SASP^5–7^. Selective elimination of senescent cells extends life-span and prevents or attenuates age-associated diseases in mice^8, 9^, and therefore, senolytic drugs have been developed^7, 10^. The primary targets of first-generation senolytic drugs are senescent cell anti-apoptotic pathways, which are upregulated in senescent cells. A combination of dasatinib (an SRC/tyrosine kinase inhibitor) and quercetin (a natural flavonoid) effectively eliminates senescent cells and reduces the senescent cell burden in model aged mice^11^. ABT-263 (Navitoclax) and ABT-737, which are pan-inhibitors of the anti-apoptotic Bcl-2 protein family members Bcl-2, Bcl-xL and Bcl-w, selectively eliminate senescent cells in different species and multiple cell types. In a mouse model, these drugs rejuvenate the stem cell population in various tissues and recover their functions^12, 13^ and attenuate the progression of age-related neurodegenerative disease by modulating senescent cells^14^. ABT-263 was also applied for anti-cancer therapy^15, 16^ and clinical trials for lymphoid malignancy and relapsed small cell lung cancer were performed, but adverse toxicity, including thrombocytopenia and neutropenia, was a concern^17, 18^.

Inhibitor of apoptosis proteins (IAPs), such as cellular inhibitor of apoptosis protein 1 (cIAP1), cellular inhibitor of apoptosis protein 2 (cIAP2), and X-linked inhibitor of apoptosis protein (XIAP), are frequently overexpressed in various human cancers and are thought to be linked to tumor progression, chemoresistance, and poor prognosis^19^. cIAPs and XIAP can directly bind to caspases via their baculovirus IAP repeat domains and downregulate caspase activity^20–22^. Second mitochondria-derived activator of caspase (SMAC, also called DIABLO), an endogenous mitochondrial protein that is released into the cytoplasm upon apoptosis induction, neutralizes the inhibitory activities of XIAP and cIAPs and promotes caspase activation^23, 24^. Various small-molecule mimetics of SMAC have been developed as IAP antagonists of both cIAPs and XIAP^25, 26^. These IAP antagonists target cIAP1 and cIAP2 for degradation via autoubiquitination, inhibit XIAP activity and induce tumor necrosis factor α (TNFα)-mediated apoptotic cell death^27–29^. Several IAP antagonists have been developed as anti-cancer therapeutic drugs either as monotherapy or combination therapy with chemotherapy and/or radiotherapy^30^. Although AT406, a potent IAP antagonist,^31^ selectively induces apoptosis of senescent bone marrow mesenchymal stem cells^32^, the senolytic effects of IAP antagonists on tumor cells treated with chemotherapeutic drugs have not been fully investigated.

In this study, we investigated the cellular response of human colorectal cancer cells that have undergone senescence upon chemotherapeutic drug treatment and the cytotoxic effects of IAP antagonists on these cells. IAP antagonists were selectively cytotoxic to TIS tumor cells that secrete TNFα via activating caspase 8. However, they were still selectively cytotoxic to TIS tumor cells even in the absence of TNFα. Furthermore, IAP antagonists sensitized non-senescent tumor cells in the presence of TNFα. These data indicate that IAP antagonists are broadly cytotoxic to tumor cells when chemotherapeutic drugs induce senescence accompanied by TNFα secretion.

## Materials and Methods

### Cell culture and reagents

HCT116 and RKO cells were obtained from the American Type Culture Collection, and confirmed to be negative for mycoplasma by MycoAlert (Lonza K.K.). All cell lines were cultured in DMEM (Nacalai Tesque) supplemented with 10% fetal bovine serum at 37°C in 5% CO2. The following reagents were used: recombinant human TNFα protein (R&D Systems); human TNFα neutralizing antibody (D1B4, Cell Signaling Technology); FTD and DOX (Tokyo Chemical Industry); CPT (Sigma); and AT406, AZD5582, ABT-199, ABT-286, S63845, A1155463, A1331852, dasatinib, quercetin, and z-VAD-fmk (Selleck).

### Cell viability assay

Cells were plated in a 96-well plate and treated with FTD, CPT, or DOX for 3 days. Cell viability was determined using a CellTiter-Glo^®^ Luminescent Cell Viability Assay (Promega) or crystal violet staining. Cell toxicity was evaluated using a CellTox^TM^-Green Cytotoxicity Assay (Promega).

### Immunoblot analysis

Cells were lysed in RIPA buffer [1.0% NP40, 50 mmol/L Tris HCl (pH 8.0), 150 mmol/L NaCl, 0.5% deoxycholate, and 0.1% SDS] supplemented with cocktails of protease and phosphatase inhibitors. Equal amounts of protein were separated by SDS-PAGE and immunoblotted. Luminescence signals were detected using an ImageQuant LAS4000 mini system (GE Healthcare). The antibodies are listed in **Supplementary Table 1**.

### Quantitative reverse transcription polymerase chain reaction (qRT-PCR)

Total RNA was extracted from cells using an RNeasy^®^ Plus Mini Kit (Qiagen) and cDNA was synthesized using SuperScript^TM^ III First-strand Synthesis Super Mix (Thermo Fisher Scientific). qRT-PCR was performed using THUNDERBIRD^®^ SYBR qPCR Mix (QPS-201, TOYOBO) on a LightCycler^®^ 480 II system (Roche Diagnostics K.K.) The sequences of primers are listed in **Supplementary Table 2**.

### SA-β-Gal staining

SA-β-Gal activity was assessed using a Senescence Detection Kit (BioVision). All images were taken on a TS100 inverted microscope with a 20× objective using a DS-Fi2 camera (Nikon). At least 200 cells were counted per sample.

### ELISA

Collected medium was concentrated using Amicon^®^ Ultra (Merck). TNFα was measured using a Quantikine Human TNFα ELISA Kit (R&D Systems). The TNFα level was normalized by the cell number at the time of cell harvesting.

### Generation of TNFα-knockout HCT116 cells

Short-guide RNA (sgRNA) sequences targeting exon 1 of the *TNFa* gene were designed using CRISPRdirect^33^. The annealed oligonucleotide containing TNFα sgRNA was cloned into the *Bbs*I site of pX330 (Addgene #42230), which was kindly provided by Dr. Feng Zhang, to generate the TNFα-sgRNA plasmid. pBluescript-G418 and VKG-sgRNA plasmids were generated as described previously^34^, and these three plasmids were co-transfected into HCT116 cells using FuGENE^®^ HD (Promega). G418-resistant clones were screened by genomic PCR and sequencing. The sequences of oligonucleotides are listed in **Supplementary Table 2.**

### Generation of conditional cIAP1- and/or XIAP-knockout cells using the mAID system

sgRNA sequences to cleave proximal to the stop codon of the *cIAP1* or *XIAP* gene were designed^33^, and the annealed oligonucleotides were cloned into the *Bbs*I site of pX330 to generate the cIAP1-sgRNA and XIAP-sgRNA plasmids. To construct donor vectors, left- and right-arm DNA fragments (350–400 bp) of each gene were amplified from genomic DNA of HCT116 cells as the template using KOD FX Neo DNA polymerase (TOYOBO) and then cloned into the *Eco*RV site of pBluescript SK+ using an In-Fusion HD Cloning Kit (Takara Bio). The *Bam*HI-digested DNA fragment containing the mAID tag and the puromycin or neomycin resistance cassette was inserted into the *Bam*HI site created between the left- and right-arm DNA fragments (**Supplementary Fig. 1A and B**). The sequences of primers and oligonucleotides are listed in **Supplementary Table 2**.

To establish cIAP1-mAID- or XIAP-mAID-knockin cells, HCT116 CMV-OsTIR1 F74G cells^35^, which were kindly provided by Dr. Masato T. Kanemaki, were co-transfected with the sgRNA plasmid of each gene and two donor DNA plasmids harboring the puromycin or neomycin resistance cassette using FuGENE^®^ HD (Promega). After double selection with puromycin and G418, colonies were screened by genomic PCR and immunoblotting. The puromycin and neomycin resistance cassettes were removed by the Cre-loxP recombination system delivered using Cre Recombinase Gesicles (Takara Bio) (**Supplementary Fig. 1A** and **B**). To establish cIAP1-mAID/XIAP-mAID-double-knockin cells, cIAP1-mAID cells, whose puromycin and neomycin resistance cassettes were removed, were co-transfected with XIAP-sgRNA plasmid and two donor DNA plasmids for XIAP-mAID harboring puromycin or neomycin resistance cassette using FuGENE^®^ HD (Promega). Immunoblot of cIAP1 and XIAP of whole cell lysates treated with an auxin analog, 5-phenyl-indole-3-acetic acid (5-Ph-IAA) (kindly provided by Dr. Masato T. Kanemaki) to confirm mAID knock-in and inducible degradation^35^.

### Statistical analysis

Statistical analysis was performed using EXSUS (CAC Croit Corp.) software.

## Results

### Senescence-inducing chemotherapeutic drugs trigger production and secretion of TNFα

Trifluridine (FTD), an active component of trifluridine/tipiracil, induces senescence of tumor cells with wild-type p53^36^. Indeed, the majority of HCT116 and RKO cells, which are colorectal cancer cell lines harboring wild-type p53, became senescence-associated β-galactosidase (SA-β-Gal)-positive when they were treated with FTD at the IC50 concentration (3 μmol/L) for 3 days (**Fig. 1A**). p53, p21, and cyclin D1 were upregulated and Rb phosphorylated at Ser 807 and 811 (pRb), cyclin B1, and Lamin B1 were downregulated (**Fig. 1B**) in these cells, indicating that FTD activated the p53-p21 signaling pathway and induced cellular senescence. Similarly, camptothecin (CPT) and doxorubicin (DOX) also induced senescence of HCT116 cells (**Supplementary Fig. 2A and B**).

**Figure 1.**
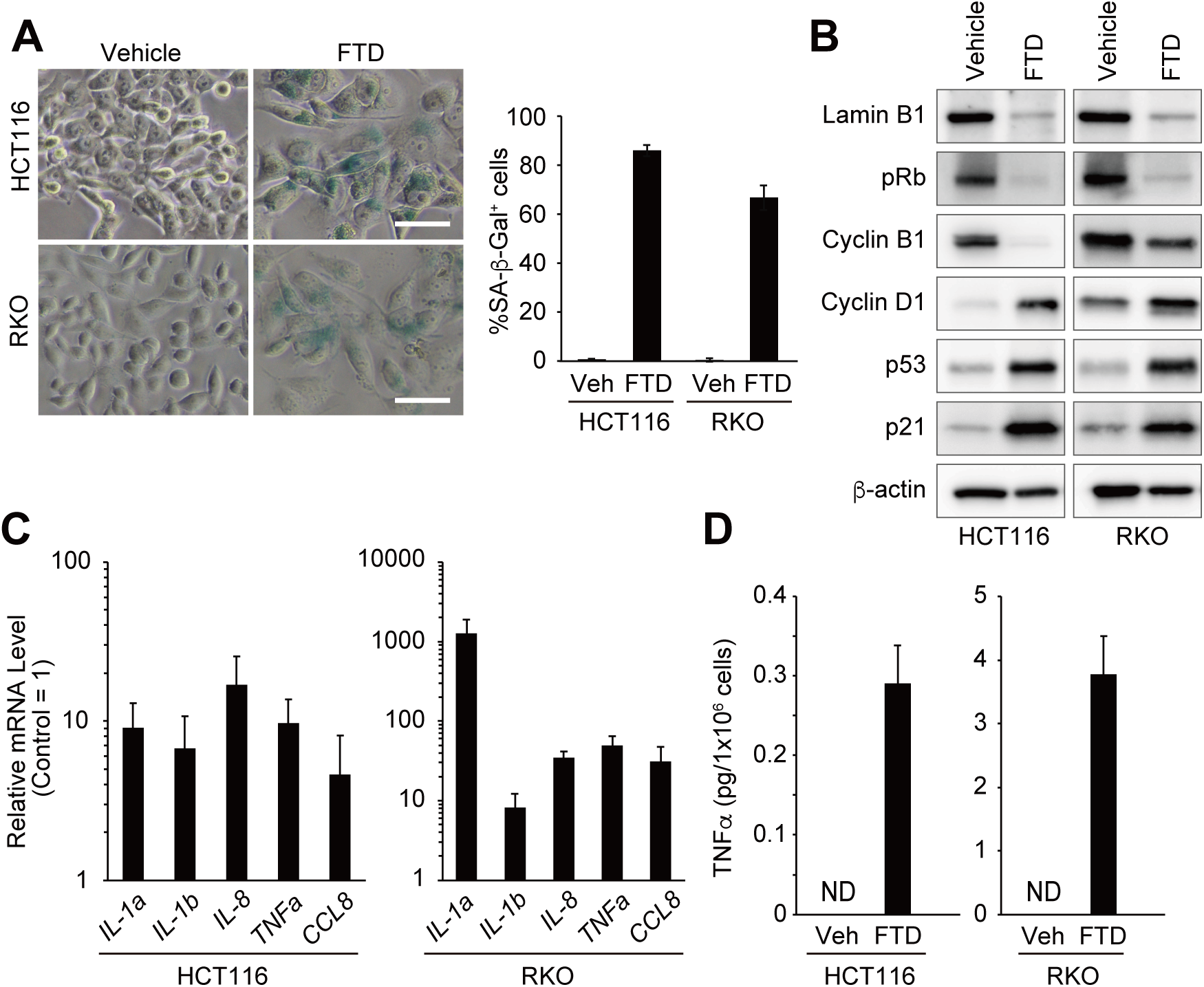
TNFα secretion by FTD-induced senescent human colorectal cancer cells. **A** SA-β-Gal activity. Cells were treated with vehicle or 3 μmol/L FTD for 3 days. The white scale bars represent 50 μm. Graphs show the percentage of SA-β-Gal positive cells. **B** Immunoblot analysis of whole cell lysates from HCT116 and RKO cells treated with vehicle or 3 μmol/L FTD for 3 days. β-actin was used as a loading control. **C** qRT-PCR analysis of senescence associated secretory phenotype (SASP) factors. Cells were treated with vehicle or 3 μmol/L FTD for 3 days. Expression of each gene was normalized by *RPLP0*. Relative mRNA level in FTD-treated cells is shown as fold change to that in vehicle-treated cells. **D** ELISA. Cells were treated with vehicle or 3 μmol/L FTD for 3 days. The amount of TNFα in the culture medium was measured. ND: not detectable. All bars and error bars represent means and standard deviation (SD), respectively, of three independent experiments.

Transcription of multiple SASP factors, including interleukin (IL)-1α, IL-1β, IL-8, TNFα, and chemokine C-C motif ligand 8 (CCL8), was increased in FTD-treated tumor cells **(Fig. 1C)**. TNFα protein was detected in the culture medium of FTD-treated tumor cells, but not in that of vehicle-treated tumor cells (vehicle, **Fig. 1D**), indicating that FTD-treated tumor cells produced and secreted TNFα. *TNFa* mRNA was also expressed in CPT- or DOX-induced senescent HCT116 cells (**Supplementary Fig. 2C**). Collectively, these results demonstrate that senescence-inducing chemotherapeutic drugs trigger production and secretion of TNFα.

### IAP antagonists selectively induce apoptosis of senescent tumor cells treated with chemotherapeutic drugs

Autocrine TNFα signaling renders human cancer cells susceptible to apoptosis induced by IAP antagonists^27–29^. To test the cytotoxic effect of IAP antagonists on senescent tumor cells, we chose AZD5582 and AT406, both of which bind to and potently antagonize XIAP, cIAP1 and cIAP2^31, 37^. Both AZD5582 and AT406 dose-dependently decreased the viability of HCT116 cells treated with FTD for 3 days, whereas their anti-proliferative effects on non-treated cells were limited (**Fig. 2A**). The selective cytotoxic effect of AZD5582 on FTD-treated HCT116 cells became more prominent as the duration of incubation increased (**Fig. 2B**). AZD5582 also dose-dependently suppressed the growth of HCT116 tumor cells treated with CPT or DOX (**Supplementary Fig. 2D**). These results indicate that IAP antagonists are selectively cytotoxic to senescent tumor cells generated following treatment with certain chemotherapeutic drugs.

**Figure 2.**
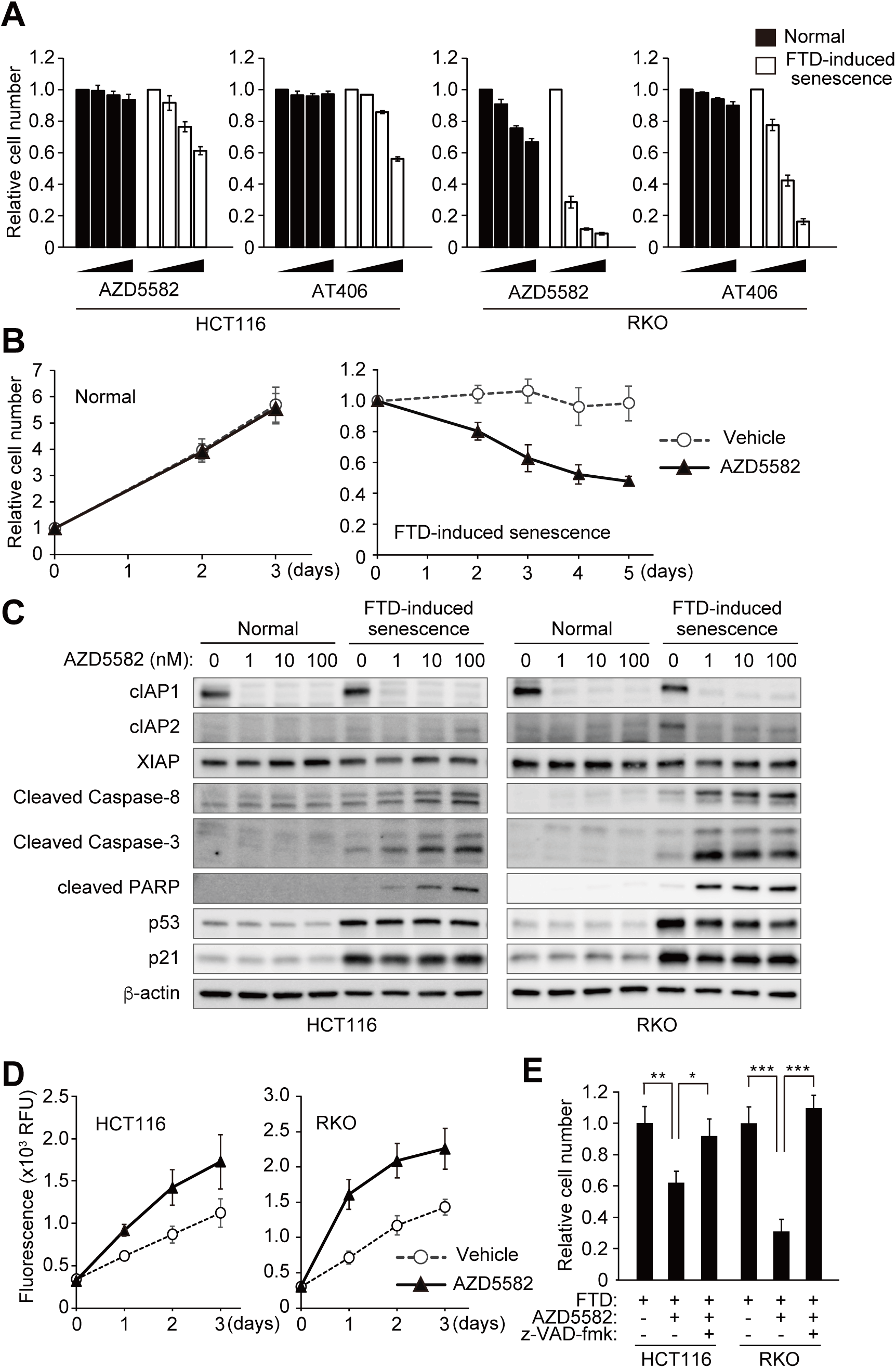
IAP antagonists selectively induces apoptosis in FTD-induced senescent tumor cells. **A** Cell viability. Growing cells cultured under normal conditions (Normal) and senescent cells induced by treatment with 3 μmol/L FTD for 3 days (FTD-induced senescence) were exposed to AZD5582 or AT406 for 3 days. Cell viability was measured by CellTiter-Glo^®^ Luminescent Cell Viability Assay. Concentrations of AZD5582 and AT406 were 0, 1, 10, 100 nmol/L and 0, 0.1, 1, 10 μmol/L, respectively. **B** Growth curve. Normal cells and FTD-treated senescent cells were exposed to vehicle or AZD5582 at 100 nmol/L. Cell viability was measured at indicated timepoint. **C** Immunoblot analysis of whole cell lysates of growing and FTD-treated HCT116 or RKO cells were incubated with AZD5582 for 24 h. β-actin was used as a loading control. **D** Cytotoxicity of FTD-treated HCT116 and RKO cells exposed to AZD5582 at 100 nmol/L Fluorescence of CellTox^TM^ Green Dye was measured at indicated timepoint. RFU: relative fluorescence unit. **E** Cell viability assay using pan-caspase inhibitor. FTD-treated HCT116 and RKO cells were pretreated with or without pan-caspase inhibitor z-VAD-fmk at 100 μmol/L for 6 h and then incubated with AZD5582 at 100 nmol/L. Cell viability was measured at 72 h (HCT116) or 24 h (RKO) after AZD5582 treatment. All bars and error bars represent means and SD, respectively, of three independent experiments. Student’s *t* test. \**p* < 0.05, \*\**p* < 0.01, \*\*\**p* < 0.001.

IAP antagonists trigger degradation of cIAP1 and cIAP2 via autoubiquitination^28, 29^. In our condition, cIAP1 was clearly degraded even when a low concentration (1 nM) of AZD5582 was added, regardless of whether tumor cells were treated with FTD (**Fig. 2C**). cIAP2 was also degraded upon treatment with 1 nM AZD5582, but was detected upon treatment with a high concentration (100 nM) of AZD5582, possibly because of the negative feedback effect of cIAP1 degradation. By contrast, the amount of XIAP was relatively unchanged. Treatment of FTD-treated tumor cells with AZD5582 induced production of the cleaved forms of PARP, caspase 3 and caspase 8 (**Fig. 2C**), indicating that the apoptosis pathway was selectively activated in FTD-induced senescent tumor cells. Consistently, AZD5582-induced cytotoxicity, as measured by fluorescent labeling of DNA in cells with impaired membrane integrity, was greater in FTD-treated tumor cells than in non-treated tumor cells (**Fig. 2D**). Furthermore, pretreatment with z-VAD-fmk, a pan-caspase inhibitor, greatly rescued AZD5582-induced growth suppression of FTD-treated tumor cells (**Fig. 2E**), confirming that IAP antagonists selectively eliminate senescent tumor cells via caspase-dependent apoptosis.

We also evaluated the cytotoxic effect of other possible senolytic drugs on HCT116 (**Supplementary Fig. 3A**) and RKO (**Supplementary Fig. 3B**) cells treated with FTD. ABT-263, which is a potent inhibitor of Bcl-xL, Bcl-w and Bcl-2^38^, A1155463 and A1331852, which are specific inhibitors of Bcl-xL^39^, and S63845, which is a specific Mcl-1 inhibitor^40^, were selectively cytotoxic to FTD-treated tumor cells. However, ABT-199, which is a selective Bcl-2 inhibitor^41^, did not exhibit such selective cytotoxicity. In our experimental setting, neither dasatinib nor quercetin^11^ showed selective cytotoxicity on tumor cells treated with FTD. These results suggest that FTD-induced senescent tumor cells become vulnerable to inhibitors of anti-apoptotic Bcl-2 family proteins, which block Bax/Bak-mediated mitochondrial outer membrane permeabilization (MOMP)^42, 43^.

### TNFα is not essential for the selective cytotoxic effects of IAP antagonists on senescent tumor cells

Caspase 8 was selectively activated by IAP antagonists in FTD-induced senescent tumor cells; therefore, TNFα secreted by FTD-treated cells may trigger extrinsic apoptotic signaling. To evaluate the contribution of TNFα to the senolytic effects of IAP antagonists, we generated *TNFa*-knockout HCT116 cells (TNFα^-/-^ cells). After FTD treatment for 3 days, SA-β-Gal-positive cells emerged to a similar level among TNFα^-/-^ cells as among parental cells (**Fig. 3A**), whereas TNFα was not detected in the culture medium (**Fig. 3B**). We evaluated the selective cytotoxic effects of IAP antagonists on FTD-treated TNFα^-/-^ cells. Intriguingly, both AZD5582 (**Fig. 3C**) and AT406 (**Supplementary Fig. 4A**) were cytotoxic to FTD-treated TNFα^-/-^ cells, whereas AZD5582 was not cytotoxic to vehicle-treated TNFα^-/-^ cells (**Supplementary Fig. 4B**), suggesting that IAP antagonists still elicit senolytic effects on FTD-induced senescent tumor cells even in the absence of TNFα.

**Figure 3.**
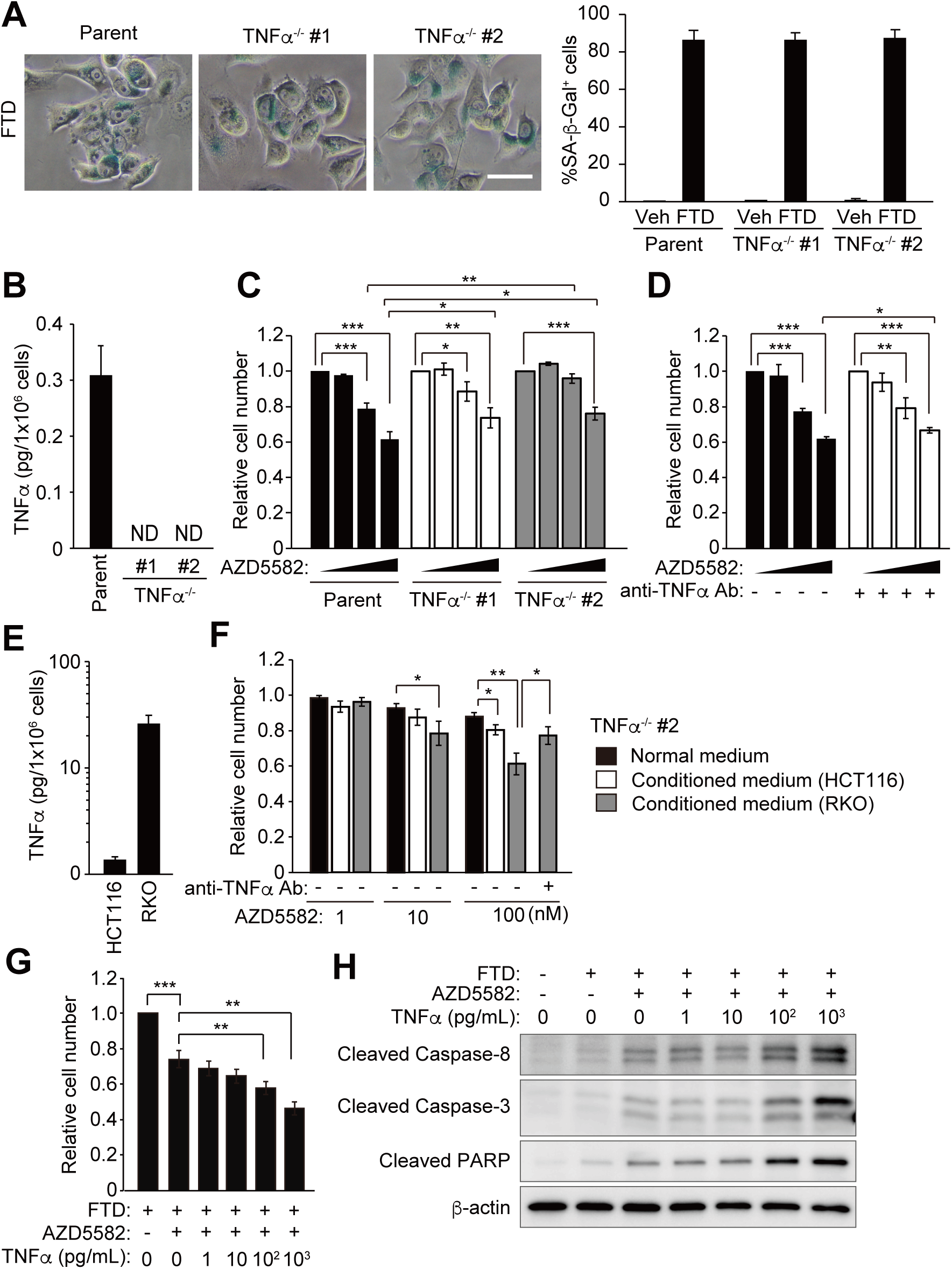
IAP antagonist eliminates senescent tumor cells in TNFα-independent manner. **A** SA-β-Gal activity of HCT116 and HCT116 TNFα^-/-^ cells treated with FTD at 3 μmol/L for 3 days. The white scale bar represents 50 μm. **B** ELISA. HCT116 and TNFα^-/-^ cells were treated with 3 μmol/L FTD for 3 days. The amount of TNFα in the culture medium was measured. ND: not detectable. **C** Cell viability of FTD-treated HCT116 and HCT116 TNFα^-/-^ cells exposed to AZD5582 for 3 days. AZD5582 concentration: 0, 1, 10, 100 nmol/L. **D** Cell viability assay using TNFα neutralizing antibody in senescent HCT116 cells. FTD-treated HCT116 cells were exposed to AZD5582 with or without TNFα neutralizing antibody for 3 days. AZD5582 concentration: 0, 1, 10, 100 nmol/L. **E** TNFα in the conditioned media. HCT116 and RKO cells were treated with 3 μmol/L FTD. After 6 days (FTD washout was conducted at day 3), conditioned media were collected. The amount of TNFα in the conditioned media was measured by ELISA. **F** Cell viability of FTD-treated HCT116 TNFα^-/-^ cells exposed to AZD5582 at indicated concentration with normal or conditioned media for 3 days. **G** Cell viability assay of FTD-treated HCT116 TNFα^-/-^ cells exposed to AZD5582 at graded doses of TNFα for 3 days. **H** Immunoblot analysis of whole cell lysates from FTD-treated HCT116 TNFα^-/-^ cells exposed to AZD5582 at graded doses of TNFα for 24 h. All bars and error bars represent means and SD, respectively, of three independent experiments. Student’s *t* test. \**p* < 0.05, \*\**p* < 0.01, \*\*\**p* < 0.001.

FTD-treated TNFα^-/-^ cells tended to be less sensitive to AZD5582 (**Fig. 3C)** and AT406 (**Supplementary Fig. 4A**) than FTD-treated HCT116 cells, but the differences were marginal. We further tested the cytotoxic effects of IAP antagonists on FTD-treated HCT116 cells in the presence or absence of a TNFα-neutralizing antibody. The TNFα-neutralizing antibody alleviated the cytotoxic effects of IAP antagonists on FTD-treated HCT116 cells, but the differences were only marginal upon treatement with 100 nmol/L AZD5582 (**Fig. 3D**) or 10 μmol/L AT406 (**Supplementary Fig. 4C**). The TNFα-neutralizing antibody did not observably alleviate the cytotoxic effects of AZD5582 (**Supplementary Fig. 4D**) or AT406 (**Supplementary Fig. 4E**) on FTD-treated RKO cells. These results indicate that TNFα does not play an essential role in the selective cytotoxic effects of IAP antagonists on FTD-induced senescent tumor cells.

We next examined whether TNFα enhances the cytotoxic effects of IAP antagonists on FTD-induced senescent tumor cells. We prepared conditioned media of HCT116 and RKO cells that had been treated with FTD for 3 days and then cultured without FTD for 3 days. The level of TNFα was approximately 10-fold higher in the conditioned medium of RKO cells than in that of HCT116 cells (**Fig. 3E**). The cytotoxic effect of AZD5582 on FTD-treated TNFα^-/-^ #2 cells was greater when the conditioned media were used, especially the conditioned medium of RKO cells (**Fig. 3F**). Furthermore, TNFα-neutralizing antibody alleviated the cytotoxic effect of AZD5582 in the conditioned medium of RKO cells (**Fig. 3F**), indicating that TNFα enhances the cytotoxic effects of IAP antagonists on senescent tumor cells. When FTD-treated TNFα^-/-^ #2 cells were exposed to AZD5582 together with purified TNFα, TNFα dose-dependently enhanced the cytotoxicity of AZD5582 (**Fig. 3G**) and upregulated the cleaved forms of caspase 8, caspase 3, and PARP (**Fig. 3H**), indicating that TNFα dose-dependently enhances the cytotoxicity of IAP antagonists by activating extrinsic caspase-mediated apoptosis.

### IAP antagonists are selectively cytotoxic to nutlin-3a-induced senescent tumor cells

TNFα, together with other SASP factors, was produced and secreted by senescent tumor cells generated following treatment with chemotherapeutic drugs (**Fig. 1C, D** and **Supplementary Fig. 2C**). The signaling pathways of the chronic DNA damage response are involved in the induction of SASP factors^44^. Nutlin-3a, which activates p53 by blocking the MDM2-p53 interaction, induces a pseudo-senescence phenotype without genotoxic stress^45, 46^. Although SA-β-Gal-positive cells emerged when HCT116 cells were treated with nutlin-3a for 3 days (**Fig. 4A**), indicators of the DNA damage response, such as activation of the ATM-Chk2 signaling pathway and γH2AX, were not detected in these cells (**Fig. 4B**). Furthermore, neither induction of *TNFa* gene expression nor secretion of TNFα into the culture medium was detected (**Fig. 4C and D**). However, AZD5582 exhibited dose-dependent cytotoxic effects on nutlin-3a-treated HCT116 cells (**Fig. 4E**), suggesting that IAP antagonists selectively eliminate senescent tumor cells in the absence of exogenous DNA damage and TNFα production.

**Figure 4.**
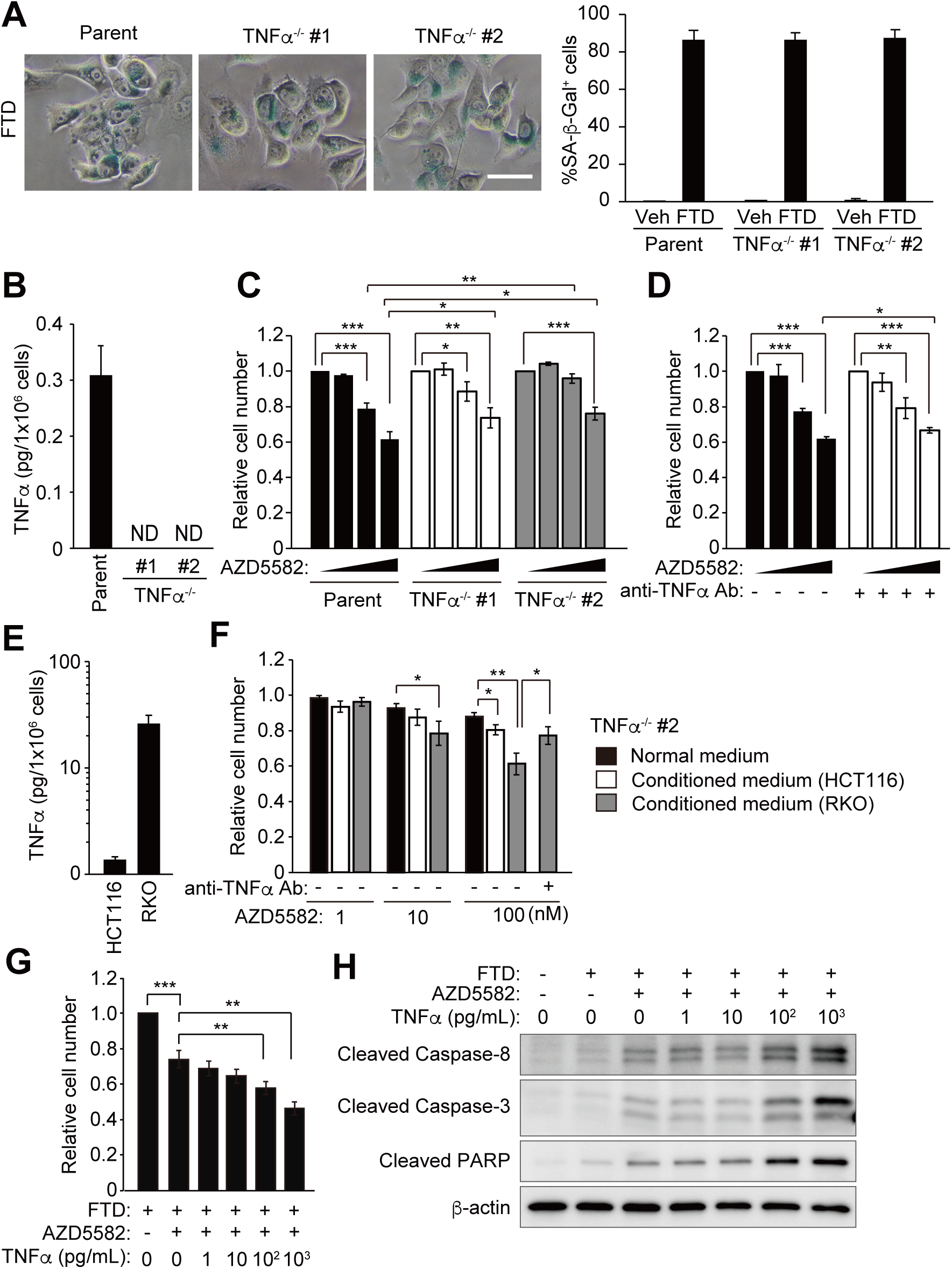
IAP antagonist confers selective cytotoxicity to senescent tumor cells without TNFα secretion. **A** SA-β-Gal activity of cells treated with vehicle or 5 μmol/L nutlin-3a for 3 days. The white scale bar represents 50 μm. Graphs show the percentage of SA-β-Gal positive cells. **B** Immunoblot analysis of whole cell lysates from HCT116 cells treated with 3 μmol/L FTD or 5 μmol/L nutlin-3a for 3 and 6 days. **C** qRT-PCR analysis of *TNFa* gene expression. HCT116 cells were treated with 3 μmol/L FTD or 5 μmol/L nutlin-3a for 3 and 6 days. Relative *TNFa* expression were shown as fold change to that in vehicle-treated cells. **D** ELISA. HCT116 cells were treated with 3 μmol/L FTD or 5 μmol/L nutlin-3a for 3 days and 6 days. The amount of TNFα in the culture medium was measured. ND: not detectable. **E** Cell viability of nutlin-3a-treated HCT116 cells exposed to AZD5582 at indicated concentrations for 3 days. **F** Cell viability of nutlin-3a-treated HCT116 cells exposed to AZD5582 at graded doses of TNFα for 3 days. **G** Immunoblot analysis of whole cell lysates from nutlin-3a-treated HCT116 cells exposed to AZD5582 at graded doses of TNFα for 24 h. All bars and error bars represent means and SD, respectively, of three independent experiments. Student’s *t* test. \**p* < 0.05, \*\**p* < 0.01, \*\*\**p* < 0.001.

To verify whether TNFα enhances the selective cytotoxic effects of IAP antagonists on nutlin-3a-induced senescent tumor cells, HCT116 cells treated with nutlin-3a and AZD5582 were exposed to various doses of TNFα. TNFα dose-dependently enhanced the cytotoxic effects of HCT116 cells treated with nutlin-3a and AZD5582 (**Fig. 4F**), and dose-dependently upregulated the cleaved forms of caspase 8, caspase 3, and PARP (**Fig. 4G**). These data indicate that exogenous TNFα enhances the cytotoxicity of IAP antagonists by activating extrinsic caspase-mediated apoptosis in senescent tumor cells without DNA damage or SASP induction.

### Suppression of both cIAP1 and XIAP is necessary for efficient cytotoxic effects on senescent tumor cells

The IAP antagonists used in this study are potent inhibitors of cIAP1, cIAP2 and XIAP^31,37^. To identify the target proteins whose inhibition is selectively cytotoxic to senescent tumor cells, we generated conditional knockdown cell lines using the modified auxin-inducible degron (mAID) system. As parental cells (Parent), HCT116 cells expressing OsTIR1 (F74G), a mutant form of the botanical E3 ubiquitin ligase for the protein with the mAID tag that is specifically activated by the auxin analog 5-phenyl-indole-3-acetic acid (5-Ph-IAA)^35^. We knocked in the mAID-encoding sequence at the 3’-end of the target gene(s) and successfully generated cells expressing cIAP1-mAID, XIAP-mAID, and cIAP1-mAID/XIAP-mAID. When 5-Ph-IAA was added to these cells, each protein was effectively degraded (**Fig. 5A**). Of note, cIAP2 protein was upregulated when cIAP1 protein was degraded, possibly because cIAP1 targets cIAP2 for degradation and cIAP1 degradation stabilizes cIAP2 protein^47, 48^.

**Figure 5.**
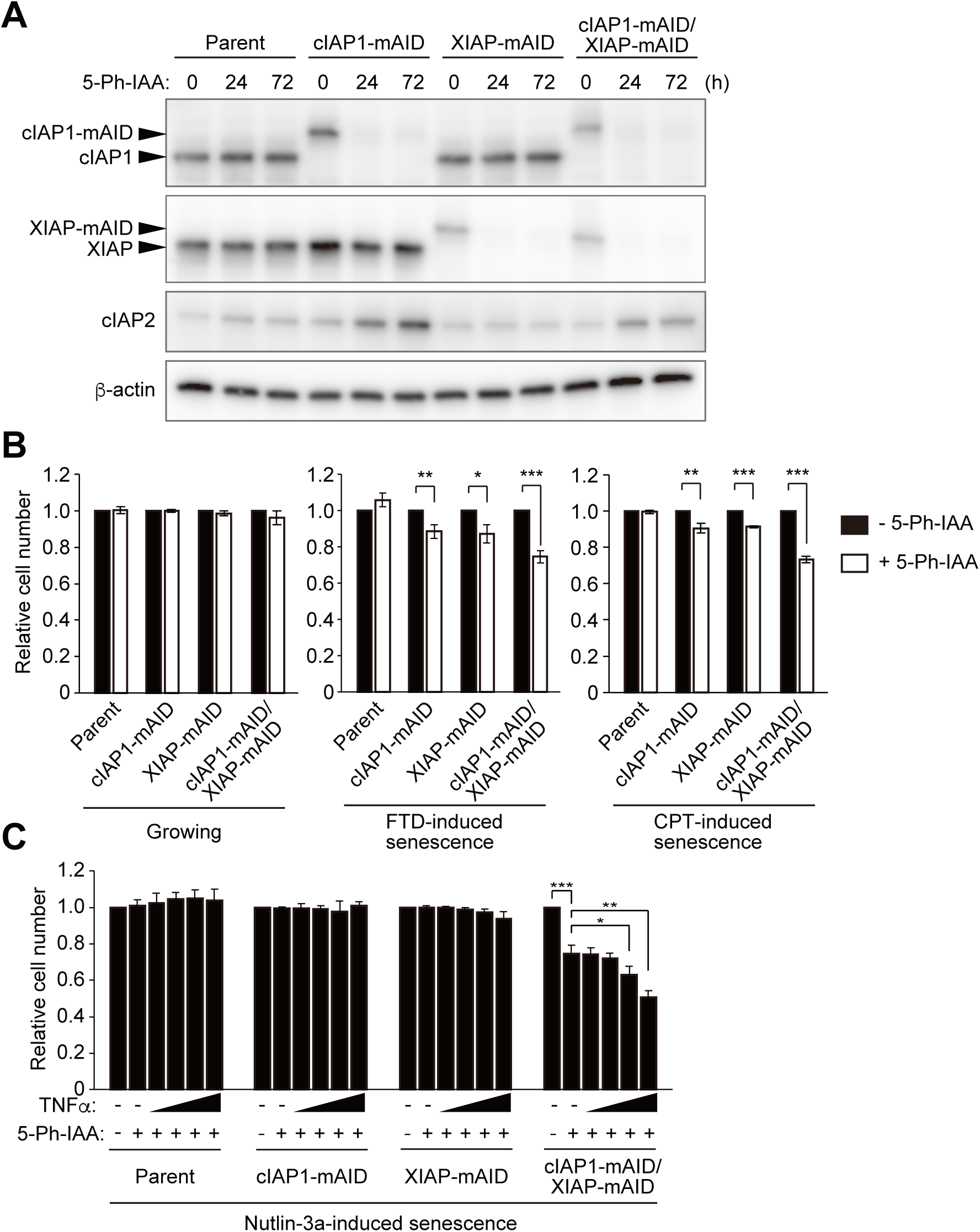
Depletion of cIAP1 and XIAP confers selective cytotoxicity to senescent tumor cells. **A** Immunoblot analysis of whole cell lysates from HCT116 cells with each auxin-inducible degron system treated with 1 μmol/L 5-Ph-IAA for 24 or 72 hrs. **B** Cell viability of growing cells cultured under normal conditions and senescent cells induced by treatment with FTD or CPT for 3 days exposed to 1 μmol/L 5-Ph-IAA for 4 days. **C** Cell viability of nutlin-3a-induced senescent HCT116 cells with auxin-inducible degron system. Nutlin-3a-treated HCT116 cells with auxin-inducible degron system were exposed to 1 μmol/L 5-Ph-IAA for 4 days with TNFα. TNFα concentration was 0, 1, 10, 100, 1000 pg/mL. All bars and error bars represent means and SD, respectively, of three independent experiments. Student’s *t* test. \**p* < 0.05, \*\**p* < 0.01, \*\*\**p* < 0.001.

In the normal growth condition, degradation of cIAP1 and/or XIAP did not affect tumor cell viability (**Fig. 5B**). By contrast, degradation of cIAP1 or XIAP reduced the number of viable FTD- or CPT-treated tumor cells, and degradation of both cIAP1 and XIAP reduced the number of viable cells more robustly than degradation of each factor individually (**Fig. 5B**), indicating these factors have an additive effect.

IAP antagonists also selectively eliminated senescent tumor cells without genotoxic stress (**Fig. 4E and F**). To identify the target proteins of IAP antagonists in this process, each mAID-knockin cell line was treated with nutlin-3a for 3 days to induce senescence and then with nutlin-3a and 5-Ph-IAA for 4 days to degrade cIAP1 and/or XIAP. Degradation of both cIAP1 and XIAP reduced the number of viable cells, but degradation of cIAP1 or XIAP did not (**Fig. 5C**). In addition, TNFα dose-dependently enhanced cytotoxicity only when both cIAP1 and XIAP were degraded (**Fig. 5C**). These results indicate that cIAP1 and XIAP have a redundant function to protect senescent tumor cells without genotoxic stress from both TNFα-independent and TNFα-induced cytotoxicity.

### IAP antagonists sensitize normally proliferating tumor cells in the presence of TNFα

Other IAP antagonists that inhibit cIAP1 and XIAP sensitize tumor cells by inducing RIPK1-Fas-associated death domain (FADD)-caspase 8-mediated apoptosis in the presence of TNFα^28, 29^. Indeed, AZD5582 dose-dependently reduced the number of normally proliferating HCT116 cells when TNFα was exogenously supplied (**Fig. 6A**). AZD5582 reduced the number of normally proliferating RKO cells even when TNFα was not supplied, and exogenous TNFα enhanced its effect (**Fig. 6B**). These results indicate that IAP antagonists induce TNFα-triggered apoptosis even in normally proliferating tumor cells unaffected by chemotherapeutic drugs.

**Figure 6.**
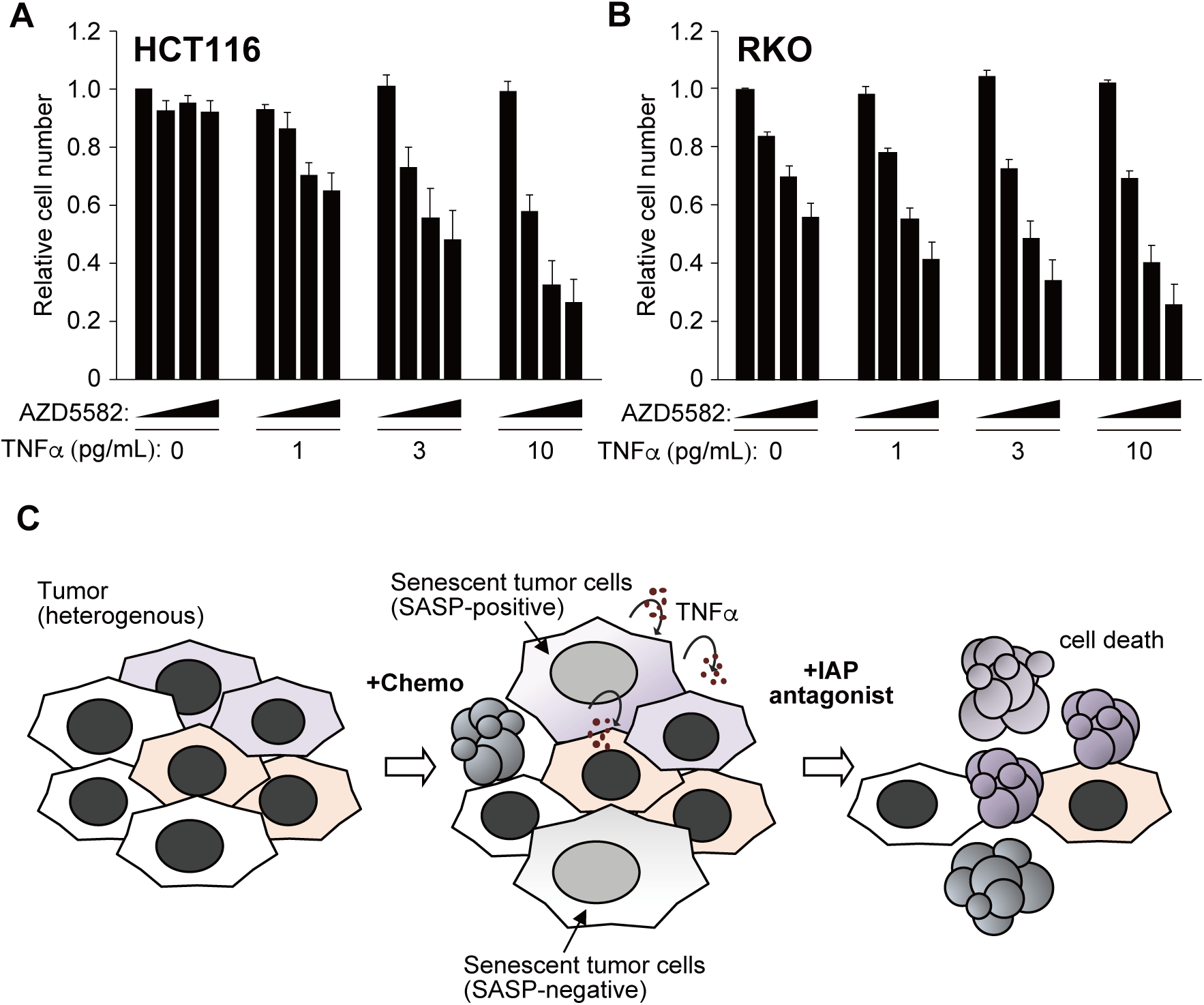
IAP antagonist and TNFα synergistically eliminate normally proliferating tumor cells A, **B** Cell viability of normally proliferating tumor cells (**A**: HCT116; **B**: RKO) treated with increasing dose of AZD5582 and TNFα. Concentrations of AZD5582 were 0, 1, 10, 100 nmol/L. All bars and error bars represent means and SD, respectively, of three independent experiments. **C** Schematic model. Tumor microenvironment is composed of heterogeneous tumor cells. When tumor is exposed to cytotoxic anti-cancer drugs (+Chemo), some tumor cells are killed, but others may fall into senescent state or keep proliferating. Some senescent tumor cells exhibit SASP-positive and secrete TNFα to the surrounding environment. If IAP antagonist is supplied, senescent tumor cells are killed irrespective of their SASP status or TNFα exposure. Furthermore, normally proliferating tumor cells under TNFα exposure are also killed by IAP antagonist. In this way, IAP antagonist will enhance the cytotoxic effect of senescence-inducing anti-cancer drug through TNFα-independent senolytic effect and TNFα-mediated cytotoxic effect.

## Discussion

TIS is a major cellular effect of anti-tumor therapy, but increases the risk of tumor dormancy and relapse^3^. To prevent future relapse, TIS tumor cells must be eradicated. In this study, we showed that TIS tumor cells secreted TNFα and became selectively sensitive to AZD5582 and AT406, which are IAP antagonists that inhibit cIAP1, cIAP2 and XIAP. These IAP antagonists induced extrinsic apoptosis via caspase 8 activation in TIS tumor cells, but not in normally proliferating tumor cells (**Fig. 2**). Intriguingly, however, selective cytotoxic effects on senescent tumor cells were still observed even in the absence of TNFα (**Fig. 3C and 4E**). Furthermore, these IAP antagonists eliminated normally proliferating tumor cells when TNFα was exogenously supplied (**Fig. 6A and B**). These data imply that IAP antagonists are good concomitant drugs of anti-tumor therapeutic drugs that induce cellular senescence because they are expected to eradicate heterogeneous tumor cells and prevent tumor relapse due to their senolytic activity and TNFα-triggered cytotoxicity.

How is TNFα produced by TIS tumor cells? The NF-κB, p38, mTOR and C/EBPβ signaling pathways are components of the SASP and are involved in chronic DNA damage response signaling^44^. *TNFa* gene expression is mainly regulated by the NF-κB pathway. DNA damage induced by chemotherapeutic drugs activates the non-canonical NF-κB pathway and consequently TNFα is produced and secreted. Upon binding of TNFα to TNF receptor I on tumor cells, the canonical NF-κB pathway is activated and TNFα is further produced. Such feedback activation of the NF-κB pathway may contribute to continuous production and secretion of TNFα. By contrast, the SASP was not induced and TNFα was not produced in senescent tumor cells generated following treatment with nutlin-3a, which activates p53 in the absence of DNA damage (**Fig. 4C and D**), possibly because the initial non-canonical NF-κB pathway was not activated.

Do IAP antagonists have senolytic activity? AZD5582 and AT406 are IAP antagonists that efficiently target cIAP1, cIAP2 and XIAP^31, 37^. Although these IAP antagonists were selectively cytotoxic to TIS tumor cells (**Fig. 2A**), this cytotoxicity included extrinsic apoptosis triggered by TNFα produced as a result of chemotherapeutic drug treatment. AZD5582 sensitized normally proliferating tumor cells in the presence of TNFα (**Fig. 6A and B**); therefore, the combined cytotoxic effect of IAP antagonists and TNFα is not senolytic. Nevertheless, our data indicate that IAP antagonists have senolytic activity because (1) AZD5582 and AT406 were selectively cytotoxic to TNFα^-/-^ TIS tumor cells (**Fig. 3C and Supplementary Fig. 4A**) and (2) AZD5582 was cytotoxic to nutlin-3a-induced senescent tumor cells (**Fig. 4E**), which did not produce TNFα (**Fig. 4C and D**). In senescent cells, MOMP occurs in a subset of mitochondria and pro-apoptotic cytochrome c is released from these mitochondria into the cytoplasm^49^. However, the amount of cytoplasmic cytochrome c is not sufficient to induce apoptosis. XIAP is a potent inhibitor of active caspase 3, 7 and 9^50–52^. cIAP1 also inhibits activation of procaspase 3 by the caspase 9-Apaf1-cytochrome c apoptosome^53^. To induce apoptosis by activating these effector caspases in senescent cells in the absence of TNFα, both cIAP1 and XIAP would have to be efficiently suppressed. Our data obtained by analyzing nutlin-3a-treated cIAP1-mAID/XIAP-mAID cells support this idea (**Fig. 5C**).

Inhibitors of anti-apoptotic Bcl-2 family proteins (Bcl-xL, Bcl-w, and Mcl-1) were selectively cytotoxic to FTD-induced TIS tumor cells (**Supplementary Fig. 3A and B**). In TIS tumor cells, these proteins suppress pro-apoptotic Bax/Bak, which induce MOMP causing release of cytochrome c and SMAC into the cytoplasm and apoptosis^42, 43^. For efficient cytotoxic effect on senescent tumor cells, targeting mitochondria using robust MOMP inducers (*i.e.* inhibitors of anti-apoptotic Bcl-2 family proteins) and compounds that mimick pro-apoptotic mitochondrial factors (*i.e.* IAP antagonists) is a promising approach. Intriguingly, a recent study showed that ABT-199 and Birinapant (TL32711), another IAP antagonist, synergize to selectively induce death of senescent cells with low toxicity to platelets^54^.

Recently, in a CRISPR screen to identify candidate genes that have senolytic potential, cellular FADD-like interleukin-1 β-converting enzyme inhibitory protein (cFLIP) was identified^55^. Suppression of cFLIP and activation of death receptor 5 by its ligand TRAIL also sensitize adjacent non-senescent cells to killing through a bystander effect mediated by secretion of cytokines^55^. Our data indicate that IAP antagonists can trigger such a bystander effect mediated via activation of TNF receptor I by its ligand TNFα, which is secreted by adjacent TIS tumor cells, in addition to their intrinsic senolytic activity (**Fig. 6C**).

Thus, IAP antagonists are expected to be developed as anti-cancer drugs. Recently, phase III clinical trials for locally advanced squamous cell carcinoma of the head and neck were performed using AT406/Debio1143/Xevinapant as combination therapies with chemoradiotherapy (NCT04458715) or intensity-modulated radiation therapy (NCT05386550)^30^. AZD5582 downregulates Mcl-1 expression in addition to suppressing cIAPs and XIAP^56^; therefore, it is expected to be clinically developed as an anti-cancer drug^57^. Several clinical trials using other IAP antagonists, such as Birinapant and Tolinapant (ASTX660), as anti-cancer drugs are ongoing^30^. IAP antagonists may open up new avenues for therapies that aim to complete cure cancer by eradicating tumor cells and preventing relapse.

## Acknowledgments

We thank Dr. Masato T. Kanemaki (National Institute of Genetics) for providing HCT116 CMV-OsTIR1 F74G cells and 5-phenyl-indole-3-acetic acid (5-Ph-IAA), and Ms. Masako Kosugi and Atsuko Yamaguchi for their expert technical assistance. We also appreciate the technical assistance from the Research Support Center, Research Center for Human Disease Modeling, Kyushu University Graduate School of Medical Sciences.

## Ethics Statement

Our study did not require ethical approval.

## Author contributions

H. Ochiiwa: Conceptualization, Validation, Formal analysis, Investigation, Resources, Data curation, Writing-original draft, Writing-Review and Editing, Visualization. T. Wakasa: Validation. Y. Kataoka: Validation. K. Ando: Validation. E. Oki: Validation. Y. Maehara: Supervision, Funding acquisition. M. Iimori: Methodology, Validation, Resources, Supervision, Funding acquisition. H. Kitao: Conceptualization, Writing-original draft, Writing-Review and Editing, Visualization, Supervision, Project administration, Funding acquisition.

## Abbreviations

5-Ph-IAA: 5-phenyl-indole-3-acetic acid
cFLIP: cellular FADD-like interleukin-1b-converting enzyme inhibitory protein
cIAP: cellular inhibitor of apoptosis protein
CPT: camptothecin; DOX, doxorubicin
FADD: Fas-associated death domain
FTD: trifluridine
IAP: inhibitor of apoptosis protein
IL: interleukin
mAID: modified auxin-inducible degron
MOMP: mitochondrial outer membrane permeabilization
qRT-PCR: quantitative reverse transcription polymerase chain reaction
SASP: senescence-associated secretory phenotype
SA-β-Gal: senescence-associated β-galactosidase
sgRNA: short-guide RNA; SMAC, second mitochondria-derived activator of caspase; TIS, therapy-induced senescence
TNFα: tumor necrosis factor α
XIAP: X-linked inhibitor of apoptosis protein.

