## Supplementary Figures&Tables for "IAP antagonists potentiate TNFα-triggered apoptosis but selectively eliminate senescent tumor cells independently of TNFα"

Supplementary Figure 1

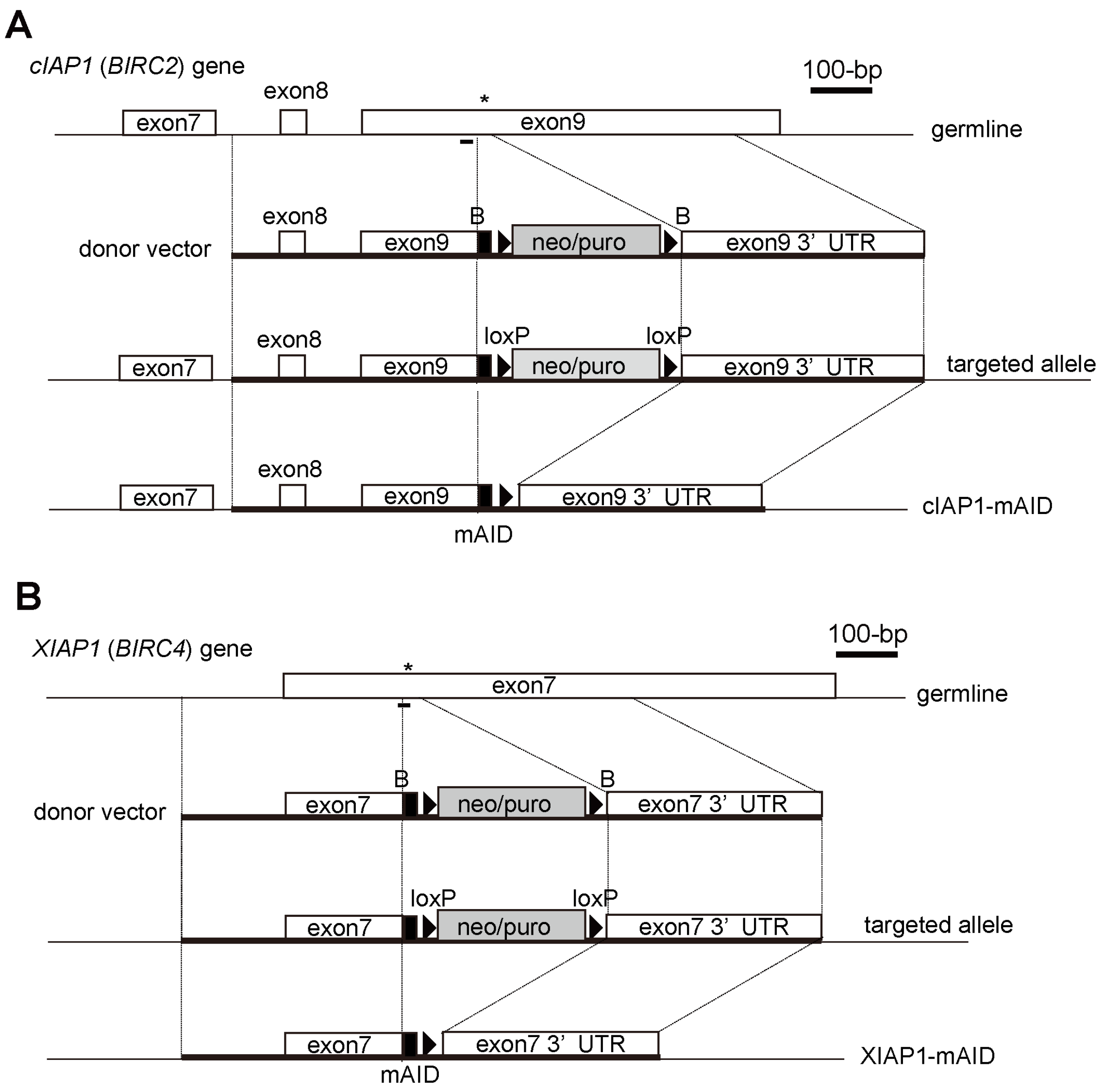


**Supplementary Figure 1**

Schematic map of cIAP1-mAID (**A**) and XIAP-mAID (**B**) knockin strategy. *: stop codon; B: *Bam*HI; Horizontal bars represent sgRNA for Cas9-mediated cleavage.

Supplementary Figure 2

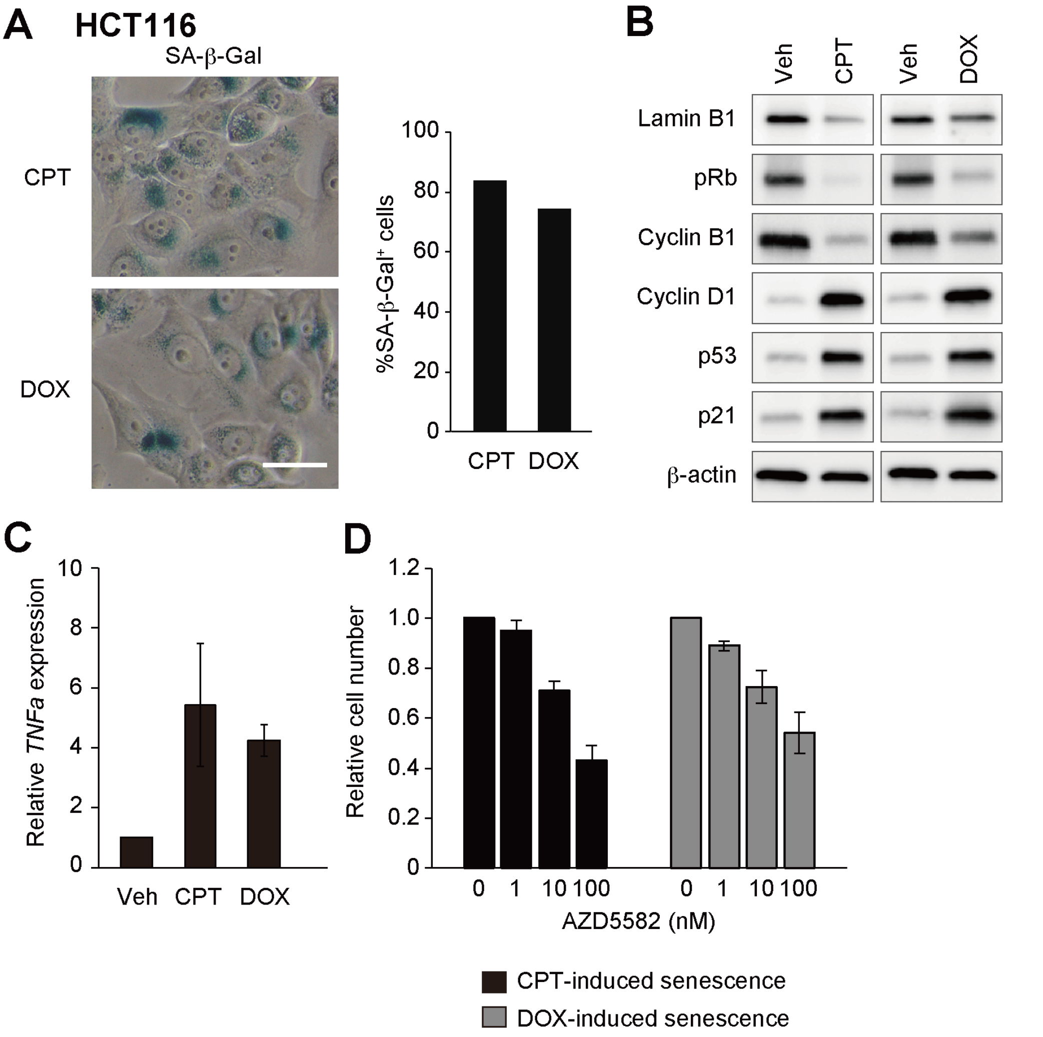


**Supplementary Figure 2**

**A** SA-β-Gal activity. Cells were treated with camptothecin (CPT) at 4 nmol/L or doxorubicin (DOX) at 200 nmol/L for 3 days. The white bar represents 50 μm. Graph shows the percentage of SA-β-Gal positive cells. **B** Immunoblot analysis of whole cell lysates from HCT116 cells treated with vehicle (Veh), 4 nmol/L CPT or 200 nmol/L DOX for 3 days. **C** qRT-PCR. HCT116 cells were treated with vehicle, 4 nmol/L CPT or 200 nmol/L DOX for 3 days. Expression levels were normalized to *RPLP0* and are shown as fold change over control. All bars and error bars represent means and SD, respectively, of three independent experiments. **D** Cell viability of senescent cells induced by treatment with 4 nmol/L camptothecin (CPT) or 200 nmol/L doxorubicin (DOX) for 3 days exposed to AZD5582 at indicated concentrations for 3 days.

### Supplementary Figure 3

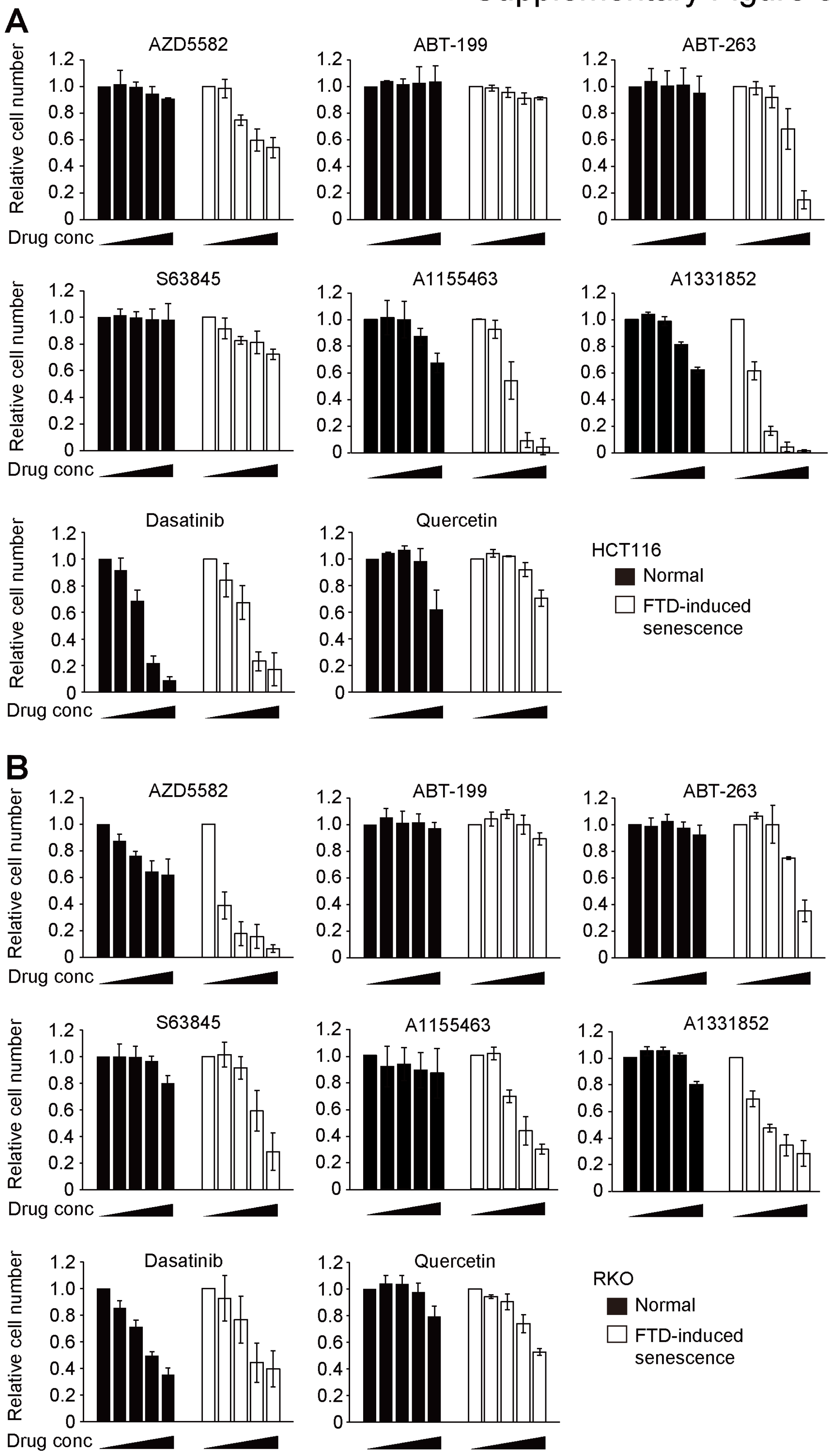


23

**Supplementary Figure 3**

Cell viability of growing cells cultured under normal condition (Normal) and senescent cells induced by treatment with 3 μmol/L FTD for 3 days (FTD-induced senescence) exposed to AZD5582 or other possible senolytic drugs for 3 days. HCT116 (**A**) and RKO (**B**) cells were used. Cell number was determined by crystal violet staining. Drug concentrations: AZD5582, ABT-199, ABT-263, A1155463, A1331852, S63845, Dasatinib, 0, 1, 10, 100, 1000 nmol/L; Quercetin, 0, 0.1, 1, 10, 30 μmol/L. All bars and error bars represent means and SD, respectively, of three independent experiments.

### Supplementary Figure 4

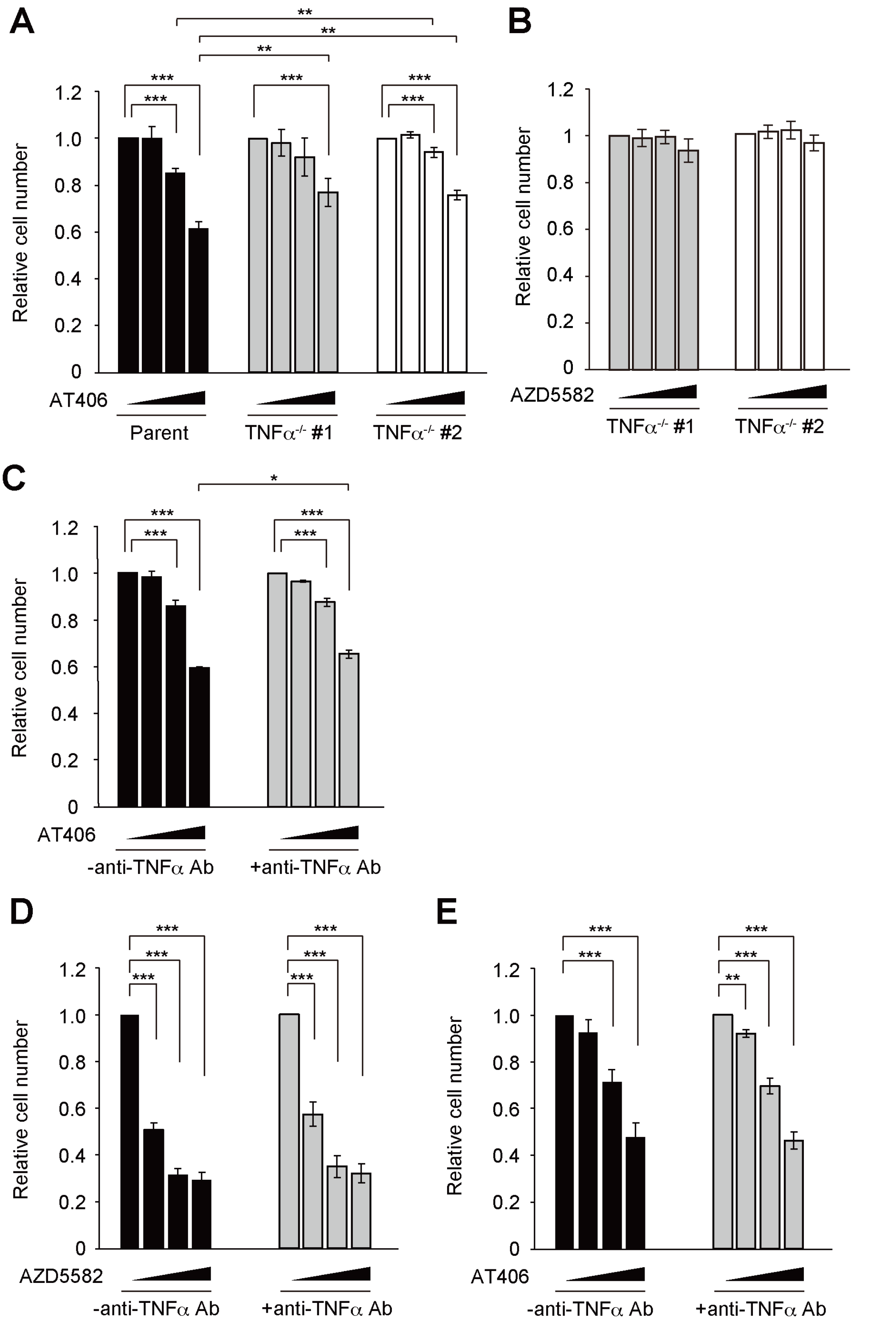

**Supplementary Figure 4**

**A** Cell viability of FTD-treated HCT116 and TNFα$^{-/-}$ cells exposed to AT406 at indicated concentrations for 3 days. AT406 concentration was 0, 0.1, 1, 10 μmol/L. **B** Cell viability of normally proliferating TNFα$^{-/-}$ cells exposed to AZD5582. AZD5582 concentration was 0, 1, 10, 100 nmol/L. **C** Cell viability of FTD-treated HCT116 cells exposed to AT406 with or without TNFα neutralizing antibody for 3 days. AT406 concentration was 0, 0.1, 1, 10 μmol/L. **D** Cell viability of FTD-treated RKO cells exposed to AZD5582 with or without TNFα neutralizing antibody for 3 days. AZD5582 concentration was 0, 1, 10, 100 nmol/L. **D** Cell viability of FTD-treated RKO cells exposed to AT406 with or without TNFα neutralizing antibody for 3 days. AT406 concentration was 0, 0.1, 1, 10 μmol/L. Cell viability was determined by CellTiter-Glo® Luminescent Cell Viability Assay.

### Supplementary Figure 5a

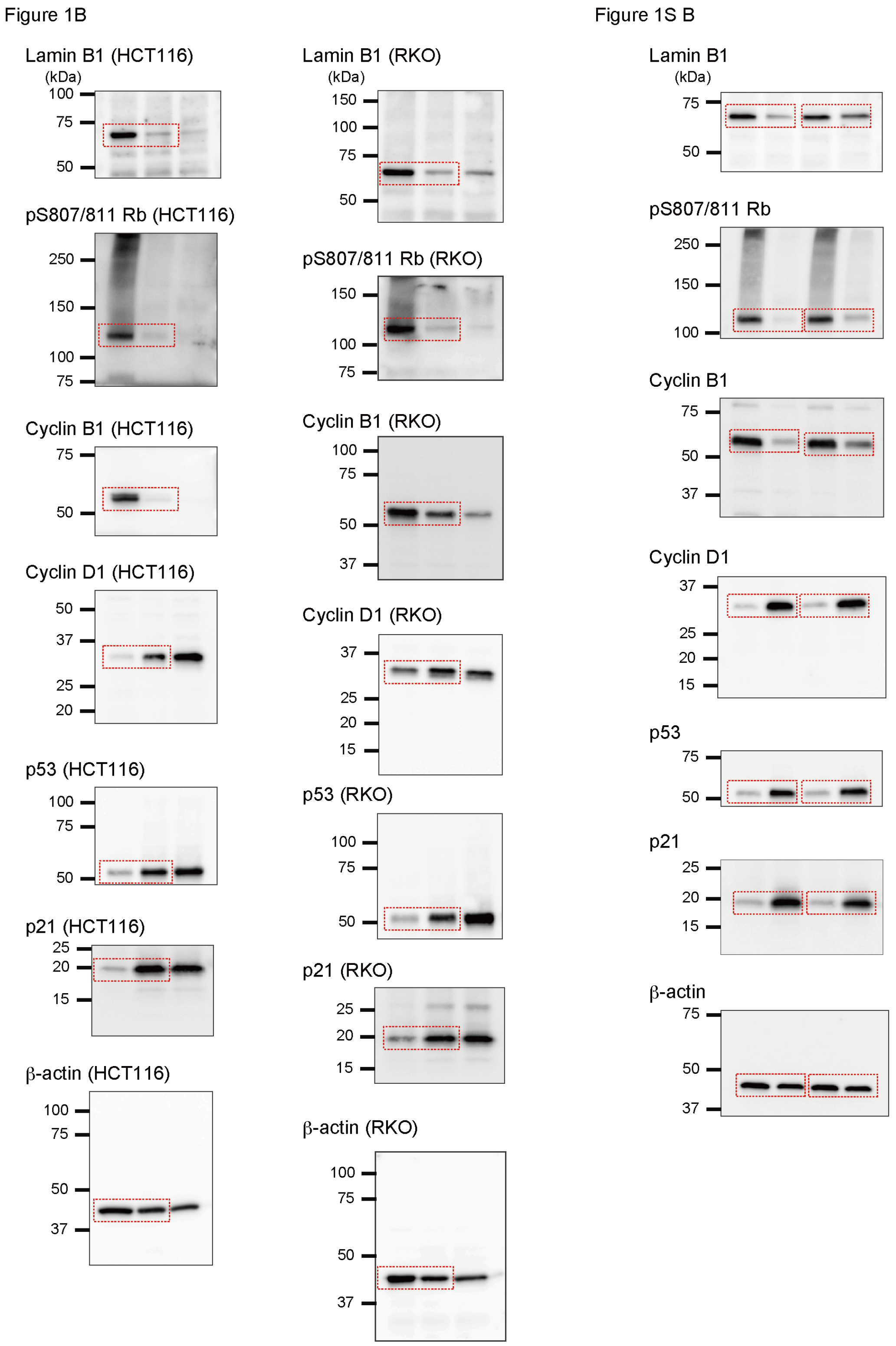

Figure 2C

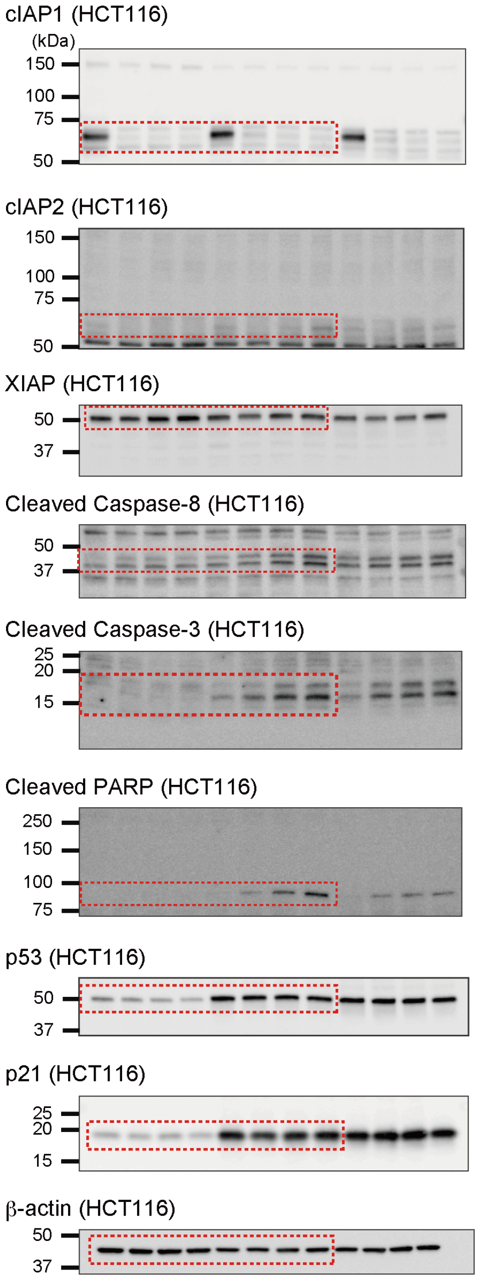


### Supplementary Figure 5b

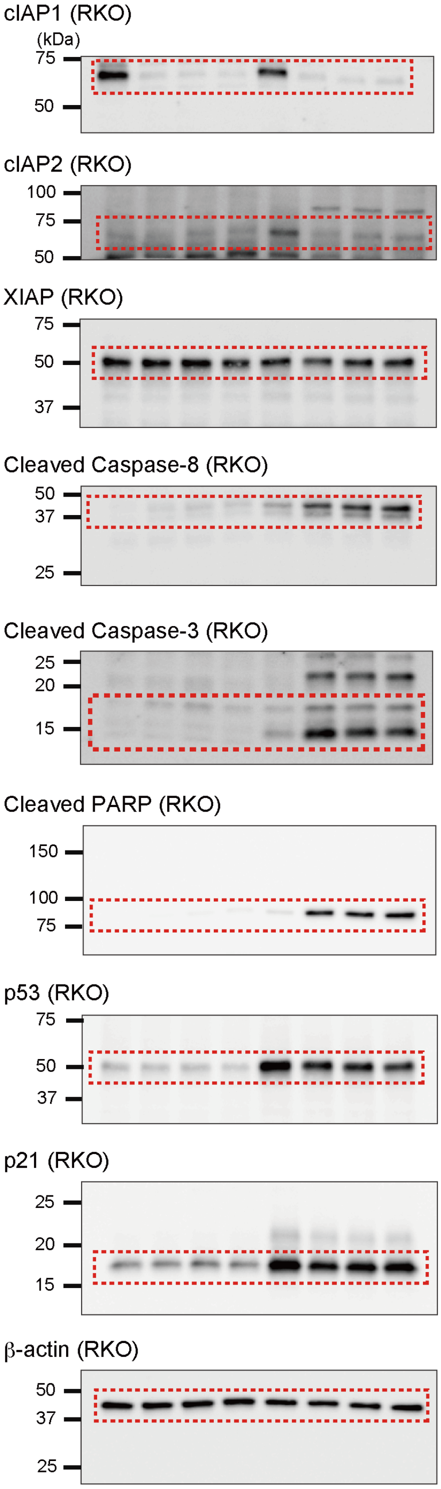

### Supplementary Figure 5c

#### Figure 3H

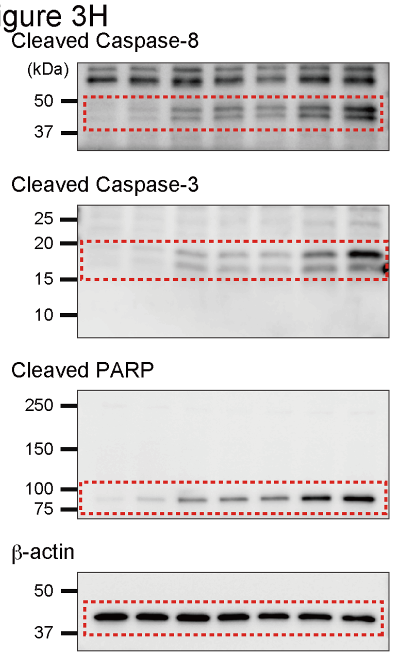


#### Figure 4B

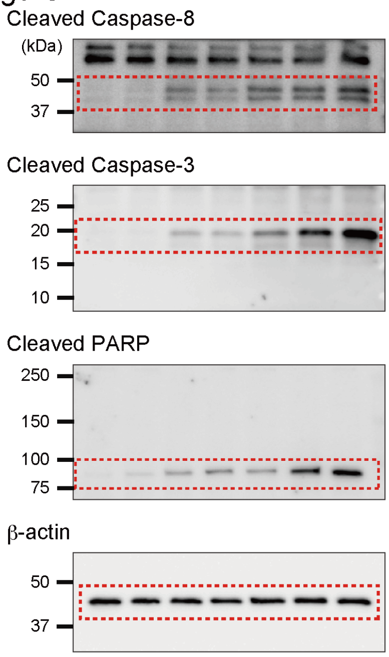


#### Figure 4G

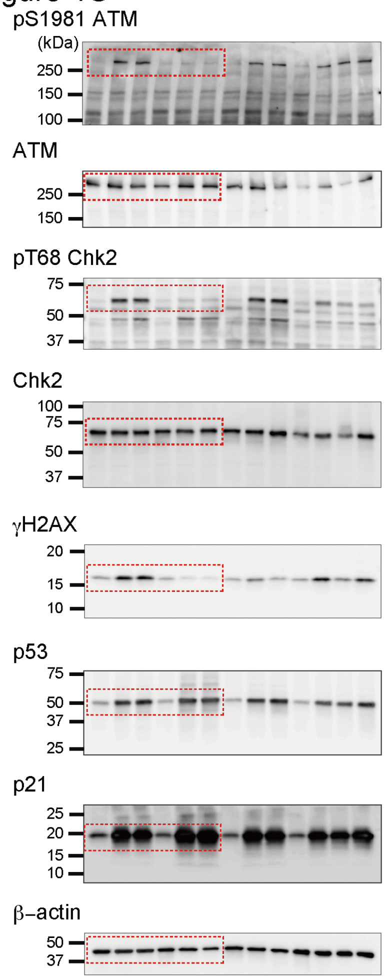


#### Figure 5A

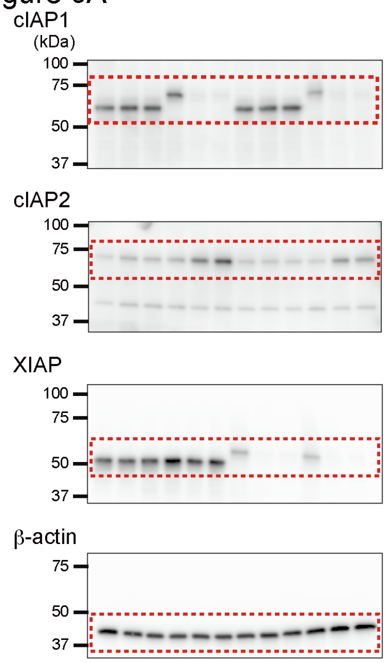

**Supplementary Figure 5**

Uncropped data of immunoblot data from Fig. 1C, Fig. 2C, Fig. 3H, Fig. 4B, 4G, and Fig. 5A.

58

Supplementary Table 1. List of antibodies

| Target protein | Supplier | Cat# |
|---|---|---|
| cIAP1 | CST | 7065S |
| cIAP2 | CST | 3130S |
| XIAP | CST | 2045S |
| Cleaved Caspase-3 | CST | 9664 |
| Cleaved Caspase-9 | CST | 7237 |
| Cleaved Caspase-8 | CST | 9496 |
| Cleaved PARP | CST | 5625 |
| LaminB1 | abcam | ab16048 |
| CyclinB1 | Millipore | 05-373 |
| p53 | DAKO | M7001 |
| p21 | CST | 2947 |
| Cyclin D1 | CST | 55506 |
| ATM | abcam | ab199726 |
| Chk2 | CST | 3440 |
| phospho ATM (Ser1981) | CST | 13050 |
| phospho Chk2 (Thr68) | CST | 2661 |
| phospho Rb (Ser807/811) | CST | 8516 |
| γH2AX | Merk Millipore | 05-636 |
| β-actin | CST | 4970 |

59

Supplemtary Table 2. List of oligonucleotides

| cIAP1-mAID knockin | Sequence |
|---|---|
| cIAP1 Left arm | F: ATCGATAAGCTTGATCAGATGCGGTGGCTCAGTTT |
| | R: TAGGATCCAGAGAGAAATGTACGAACAGTACCTTTAATGATTCCTCTGCAAATAGG |
| cIAP1 Right arm | F: ATTTCTCTCTGGATCCTAGTCTATATTTTAACCTGC |
| | R: CTGCAGGAATTCGATAAGCACCAAAGACAATTCGGC |
| cIAP1-sgRNA | S CACCTATTTGCAGGGGTATAATCA |
| | AS: AAACTGATTATACCCCTGCAAATA |
| XIAP-mAID knockin | |
| XIAP Left arm | F: ATCGATAAGCTTGATCCACTAGCGTGTGAGCTATT |
| | R: ACATAACATGGGATCCAGACATAAAAATTTTTTGCT |
| XIAP Right arm | F: CTGGATCCCATGTTATGTTGTTCTTATTACC |
| | R: CTGCAGGAATTCGATCAGAAAGCTCCATTTGTTAAGCCT |
| XIAP-sgRNA | S: CACCTCTTAATCTAACTCTATAGT |
| | AS: AAACCGTCCCGGATCATGCTTTCA |
| TNFα KO | |
| TNFα-sgRNA | S: CACCTGAAAGCATGATCCGGGACG |
| | AS: AAACCGTCCCGGATCATGCTTTCA |
| VIKING method | |
| VKG-sgRNA | S: CACCGTCGCTGCGCTCGGTCGTT |
| | AS: AAACAACGACCGAGCGCAGCGAC |
| qPCR | |
| IL8 | F : ACAAACTTTCAGAGACAGCA |
| | R : TACACACAGTGAGAATGGTTCC |
| CCL8 | F : TCACCTGCTGCTTTAACGTG |
| | R : CACAGCTTCCTTGGGACATT |
| IL-1α | F : GAAGAGACGGTTGAGTTTAAGCC |
| | R : CAGGAAGCTAAAAGGTGCTGA |
| TNFα | F : CTGCACTTTGGAGTGATCGG |
| | R : GGGTTTGCTACAACATGGGC |
| RPLP0 | F : TTCGACAATGGCAGCATCTACAA |
| | R : CTGCAGACAGACACTGGCAACA |
| IL-1β | F : CAGCTACGAATCTCCGACCAC |
| | R : GGCAGGGAACCAGCATCTTC |
